# Activating ligands of Uncoupling protein 1 identified by rapid membrane protein thermostability shift analysis

**DOI:** 10.1101/2022.02.03.478984

**Authors:** Riccardo Cavalieri, Marlou Klein Hazebroek, Camila A. Cotrim, Yang Lee, Edmund R. S. Kunji, Martin Jastroch, Susanne Keipert, Paul G. Crichton

## Abstract

Uncoupling protein 1 (UCP1) catalyzes mitochondrial proton leak in brown adipose tissue for heat production, and may combat metabolic disease if activated in humans. During the adrenergic stimulation of brown adipocytes, free fatty acids generated from lipolysis activate UCP1 via an unclear interaction. Here, we have utilized membrane protein thermostability shift analysis to characterize the interaction of activating molecules with purified UCP1. We reveal that activators influence the protein through a specific destabilizing interaction, behaving as transport substrates that shift UCP1 to a less stable conformation of a transport cycle. Through the detection of specific stability shifts in screens, we identify novel activators, including the drug ibuprofen, where ligand analysis indicates a relatively wide structural specificity for interacting molecules. Ibuprofen induces UCP1 activity in liposomes and isolated brown fat mitochondria, but not in cultured brown adipocytes. Though the drug does induce activity in UCP1-expressing HEK293 cells, demonstrating that the targeting of UCP1 in cells by approved drugs is in principle achievable as a therapeutic avenue, but requires variants with more effective delivery in brown adipocytes.

## Introduction

Brown adipose tissue of mammals has the specialized ability to oxidise nutrients to generate heat for defence of body temperature against the cold (1), and may help combat metabolic disease in humans. Brown fat occurrence in adults correlates with leanness (2–4) where activation of the tissue increases energy expenditure, lipid turnover, glucose disposal and insulin sensitivity, consistent with a positive impact on systemic metabolism and health (5–8). Thermogenesis by brown fat is achieved through the activation of Uncoupling protein 1 (UCP1), a 33 kDa mitochondrial carrier that catalyses the leak of protons across the mitochondrial inner membrane, dissipating the protonmotive force to release energy as heat at the expense of ATP synthesis (see (9) for review). Following the adrenergic stimulation of brown adipocytes, e.g. by norepinephrine following cold exposure, intracellular cAMP-dependent signalling initiates lipolysis and the release of long chain fatty acids, which directly interact to activate UCP1, overcoming the inhibition of the protein by cytosolic purine nucleotides (10). Methods to encourage brown fat development and the activation of UCP1 in the absence of physiological stimuli is a therapeutic strategy to combat diabetes, obesity and related diseases, e.g., by pharmacological activation of adipocyte β_3_-adrenergic receptors (6, 11).

How fatty acids induce proton leak by UCP1 is debated (see (9) for biochemical models). Fatty acid anions may act as a co-factor to complete a proton transport channel within the protein (12) or, instead, act as a transport substrate that is exported by UCP1 to flip back directly across the mitochondrial inner membrane in a protonated form, independent of the protein, to give a net proton transfer (13, 14). Alternatively, fatty acids in both protonated and deprotonated forms may be transported by UCP1, where long chain species remain bound to the protein following transport across the membrane, allowing just protons to be chaperoned down their electrochemical gradient (15). The fatty acids may also compete directly or act allosterically to remove inhibitory purine nucleotides, which bind to UCP1 from the cytosolic side (16).

UCP1 is a member of the mitochondrial carrier family of metabolite transporters, which all share the same basic fold and membrane topology. They are comprised of three ~100 amino acid repeat domains, each composed of two transmembrane α-helices linked by a matrix loop and small α-helix, which together form a six transmembrane helix barrel arrangement with three-fold pseudosymmetry (17, 18). Our purification studies of native UCP1 have revealed that the protein is a monomer and tightly binds three cardiolipin molecules to maintain stability, similar to other mitochondrial carriers (19). Though to what degree UCP1 employs a common carrier metabolite transport mechanism, as recently resolved for the ADP/ATP carrier (20), for proton leak is not clear. The protein has been postulated to employ a mechanistically distinct process for proton leak (21). UCP1 has been studied for many years, yet the location and molecular nature of the fatty acid binding site has not been clarified. For membrane proteins in general, there has been a lack of efficient methods to report on ligand interactions due to the practical complications imposed by membrane hydrophobicity and common need for detergents. Fatty acids add further challenges as ionic surfactants, binding non-specifically with potential to denature proteins. Information on fatty acid activator binding to UCP1 has largely been indirect and inferred from changes in nucleotide binding or activity measurements (e.g. (16, 22, 23)). An improved understanding of the activator interaction may provide therapeutic avenues to target UCP1 directly for activation to engage thermogenesis.

Here, we demonstrate the efficient detection of activator interactions with purified UCP1 through protein thermal stability shift analysis. The approach not only informs on the nature of the interaction process but provides an effective screening avenue to identify novel molecules with potential to activate UCP1 in cells.

## Results

### UCP1 thermostability shifts detect a specific destabilizing interaction with activators

We have applied a rapid fluorescence-based assay to assess the relative thermostability of purified UCP1, which can provide practical information on membrane protein integrity and interaction with detergent, lipid and ligands (24). Samples in the assay are subjected to a temperature ramp while protein unfolding is monitored by an increase in fluorescence of the probe *N*-[4-(7-diethylamino-4-methyl-3-coumarinyl)phenyl]maleimide (CPM), which reacts with protein thiols as they become solvent exposed due to denaturation to give a fluorescent adduct (cf. (25)). When assessed, UCP1 purified in dodecyl maltose neopentyl glycol (12MNG) detergent exhibited a background fluorescence consistent with at least one solvent-exposed cysteine residue present in the native state, with a transition to a higher plateau as remaining buried cysteines are revealed during protein unfolding (Fig. 1A, *top*), as observed previously in similar detergents (19, 24). Our sequencing of the ovine UCP1 gene coding region verified the presence of 8 cysteine residues in the protein (Fig. EV1; rather than 9 reported in wider sequencing data elsewhere, e.g. (26)). The peak in the derivative of the unfolding transition provides an apparent ‘melt temperature’ (*T*_m_) as a relative measure of thermal stability for a given protein population (24) (e.g. 51.0 °C for UCP1, Fig. 1A, *bottom*). This measure relates to the strength and sum of the molecular bonding that occurs within and between associated components, which contribute to the overall protein stability. Accordingly, we have found the apparent *T*_m_ of membrane proteins to exhibit distinct shifts (Δ*T*_m_) in the presence of ligands, consistent with reporting on the bonding changes associated with ligand interaction (19, 24, 27).

**Figure 1.**
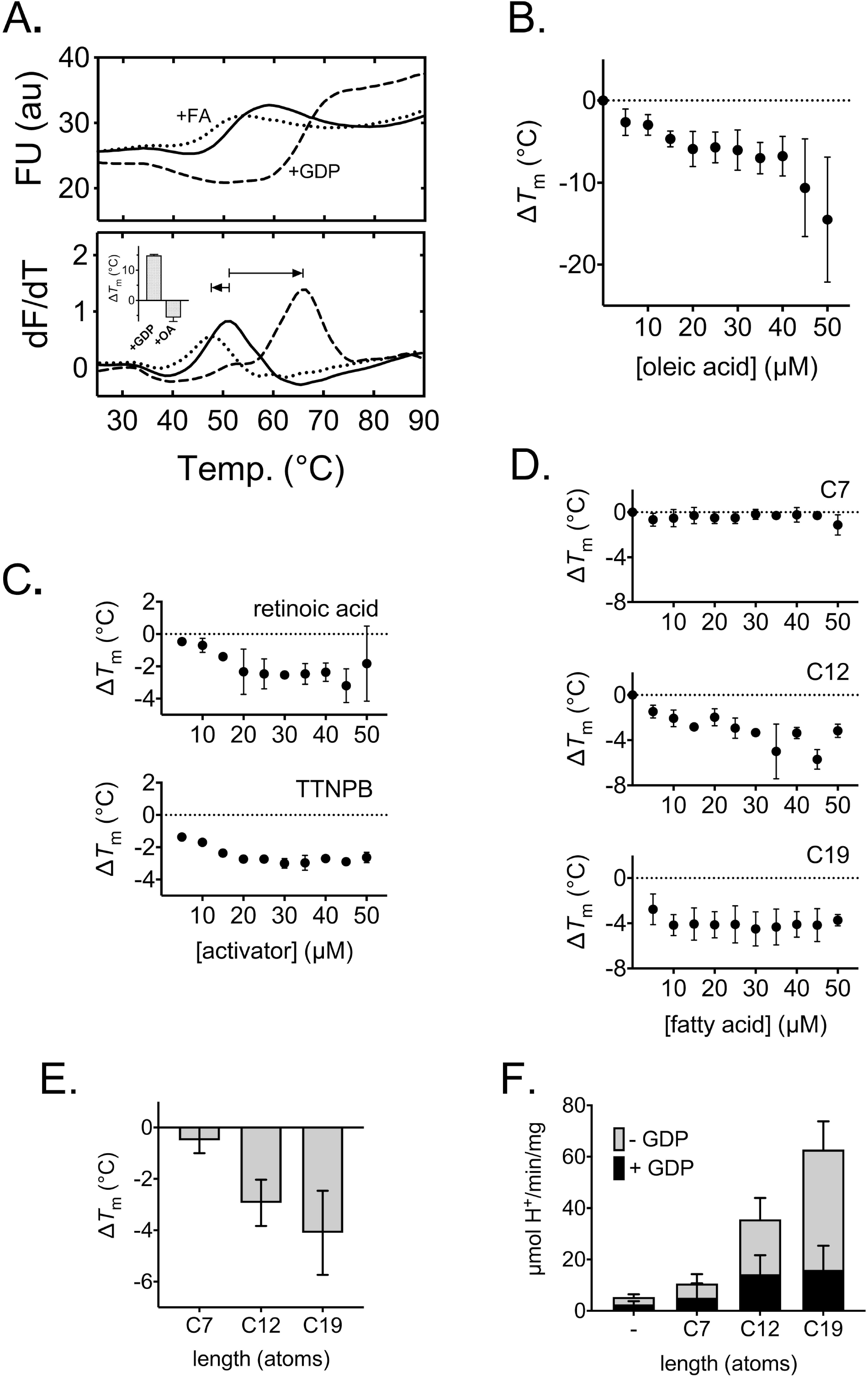
Activators induce a specific destabilization of native UCP1. The relative thermal stability of purified UCP1 monitored by the fluorescence of CPM-adduct formation at cysteine residues as they become solvent-exposed due to thermal denaturation (see Methods). (A) The thermal denaturation profile (top), and corresponding first derivative (bottom), of native UCP1 in assay buffer with 0.1%12MNG detergent in the absence (solid line) or presence of 1 mM GDP (dashed line) or 25 μM oleic acid (dotted line). *Inset*, average (± SD) shift in UCP1 ‘melt temperature’ (Δ*T*_m_; condition minus control) associated with either ligand. (B-D) Corresponding shifts in UCP1 thermal stability associated with the addition of the indicated concentration of oleic acid (B), retinoid activators (C) or saturated fatty acids of differing carbon alkyl chain length (D, and E, at 25μM specifically; C7, heptanoic acid; C12, dodecanoic acid; C19, nonadecanoic acid). (F) The corresponding proton uptake rates (nmol H^+/^min/mg protein) by UCP1 reconstituted into liposomes in the absence or presence of 100 μM C7, C12 or C19, with and without 1 mM GDP, as indicated. Data are averages (±SD) of 3-4 independent experiments for Δ*T*_m_ values and 4 independent experiments for proton uptake rates.

GDP increases the stability of UCP1 (e.g. with a Δ*T*_m_ of +15°C at 1 mM, Fig1A, *bottom*), where larger stabilisation is observed in conditions where the inhibitor is known to bind more tightly (cf. (24)). In contrast, when tested, we found that the UCP1 activator oleic acid *destabilized* UCP1 (Δ*T*_m_ of −6 °C at 25 μM, Fig. 1A, *bottom*). Fatty acids can act as harsh ionic soap at neutral pH and potentially denature proteins non-specifically. Importantly, however, when the oleic acid concentration was varied, the drop in *T*_m_ did not magnify as a simple function of the concentration, as would be expected of a non-specific effect, but instead plateaued towards a Δ*T*_m_ of −6 °C between 20 and 40 μM oleic acid (Fig. 1B), consistent with saturation of a specific interaction event that results in net decreased bonding within the UCP1 population. Above ~40 μM, a second less consistent phase was observed with an increased drop in stability with oleic acid concentration, likely representing the saturation of the species in the protein-detergent micelles and the onset of generic protein denaturation.

Further tests with other known activators of UCP1 revealed similar trends, consistently showing a destabilizing effect. The UCP1 activators retinoic acid and TTNPB (cf. (28, 29)) both destabilized UCP1 with a saturating behaviour when assessed over similar concentrations (towards a maximum Δ*T*_m_ of ~ −3 °C, Fig. 1C), as did the long chain fatty acid nonadecanoic acid (C19; towards a Δ*T*_m_ ~ −4 °C, Fig 1D). The shorter chain and less potent activator of UCP1 dodecanoic acid (C12) also destabilized UCP1 but with smaller Δ*T*_m_ shifts and without a distinguishable saturation profile, whereas the shortest fatty acid tested, heptanoic acid (C7), showed almost no destabilization at all (Fig. 1D and E). Notably, the size of the shift observed had a strong correlation with fatty acid length and ability to activate UCP1, as verified by proton leak activity assays with UCP1 proteoliposomes (cf. Fig. 1F and 1E). These thermostability measurements reveal that a specific destabilization of purified UCP1 is fundamental to the interaction with activators, and suggests a net reduction in total bonding occurs in the protein population relating to the activation process.

### Activators interact with UCP1 as transport substrates

Ligand-protein binding generates new bonds, hence, an apparent decrease in overall bonding suggests significant conformational changes occur elsewhere in the protein to account for bond loss, such as those that occur in transporter proteins to facilitate substrate translocation (cf. (30)). UCP1 has been proposed to transport fatty acids in some mechanistic models of proton leak (see above), which is supported by the observed transport of the fatty acid analogues alkyl sulfonates by the protein (13, 15, 31). In contrast to fatty acids, these species cannot be protonated at physiological pH, having very low pKa values (~ −2), and retain a negative charge that prevents them from diffusing directly across lipid membranes. These anions are transported by UCP1 but, importantly, they do not activate proton leak by the protein (13, 15, 31). When tested, we found that they induced almost identical trends in UCP1 thermostability to their equivalent fatty acid counterparts. Similar to C19, octadecane sulfonate (C18-S) destabilized the protein, saturating towards a Δ*T*_m_ of ~ −6 °C between 20 and 40 μM (Fig. 2A). Likewise, the shorter chain variants, undecane sulfonate (C11-S) and hexane sulfonate (C6-S) showed smaller shifts, matching C12 and C7, respectively, where the overall amplitude of the change observed correlated with chain length (Fig. 2A and B). In further tests with an alternative long chain fatty acid analogue, oleoyl-L-α-lysophosphatidic acid (OLPA), an inhibitor of UCP1 activity (15)), only minor shifts were observed in contrast to C18-S and C19 (Fig. 2C). These correlations indicate that activators destabilize UCP1 by interacting in an identical manner to transport substrates, rather than as activators of proton conductance *per se*, and that fatty acid anion transport is fundamental to the activation process. Of significance, the related ADP/ATP carrier was also found to be destabilized by micromolar concentrations of transport substrate (ADP) in similar assay conditions (24), suggesting a common substrate interaction process in both of these carriers.

**Figure 2.**
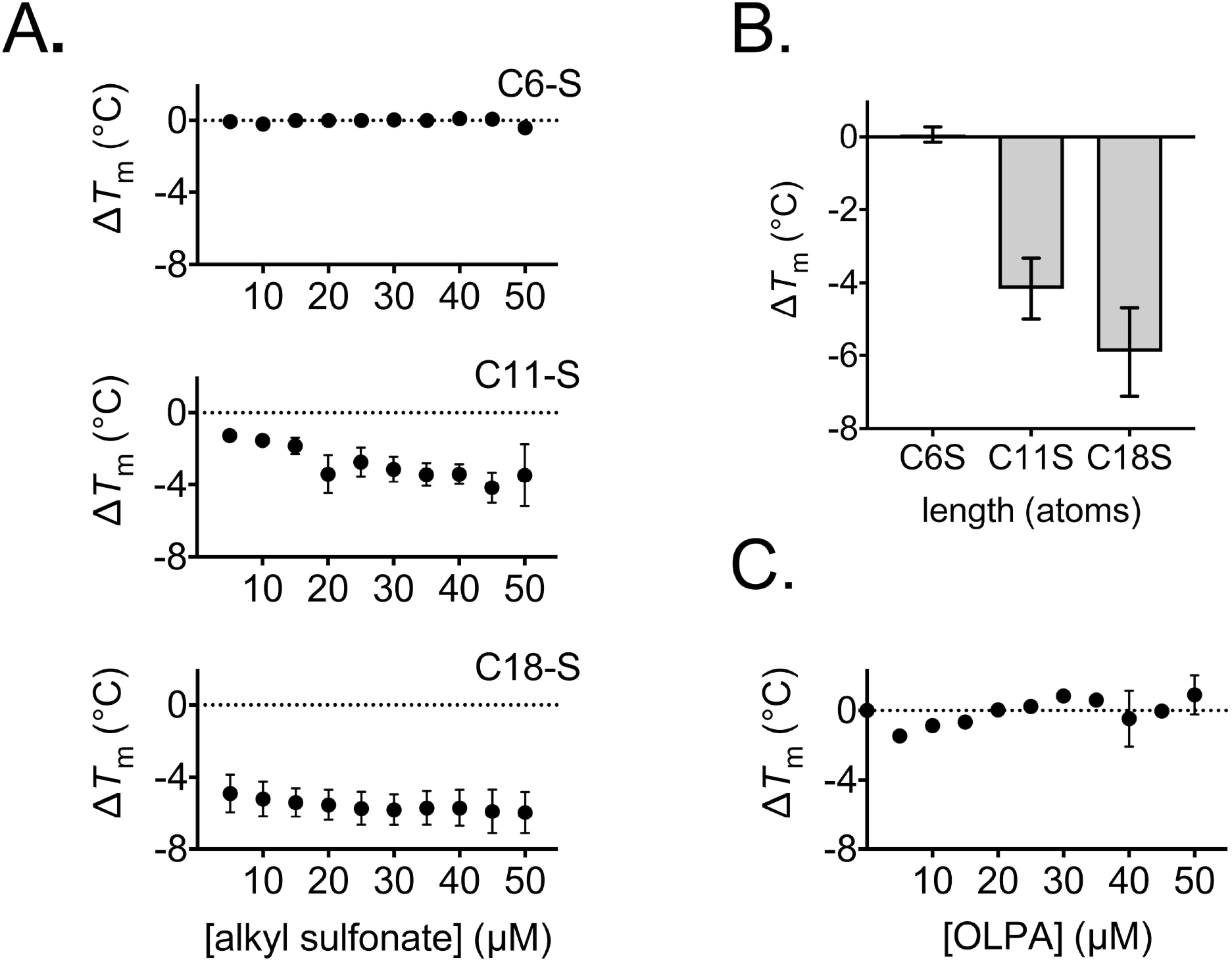
Alkyl sulfonate transport substrates and fatty acids induce the same specific destabilization of native UCP1. The relative thermal stability of purified UCP1 was determined as described for Fig. 1 (see Methods). The shifts in UCP1 thermal stability (Δ*T*_m_) associated with the addition of the indicated concentration of alkyl sulfonates (A) (C6-S, hexane sulfonate; C11-S, undecane sulfonate; C18-S, octadecane sulfonate; equivalent in atom length to C7-C19, cf. Fig. 1), or at 25 μM specifically (B), or with the addition of OLPA inhibitor (C), as indicated. Note, C18-S tests included 1.6 mM methyl-β-cyclodextrin solubilizing agent. All values given are averages (±SD) of 3 independent experiments.

### Novel activators of proton leak are found among compounds that destabilize UCP1

In light of the distinct stability changes detected with UCP1 activators, we used thermostability shift assays to screen a further 72 selected compounds, including fatty acid analogues, metabolites and drugs, to identify interacting species of potential significance (Fig. EV2). This approach identified 20 compounds that induced a significant Δ*T*_m_ in UCP1, 16 that were destabilizing and four that were stabilizing (Fig. 3A). Nine of the compounds that destabilized have been reported previously as activating ligands of proton leak or anions transported by UCP1 (15, 22, 32, 33), or were drugs of the retinoid class (adapalene, acitretin and tazarotene) related to the known UCP1 activator retinoic acid (29). Whereas one of the compounds that stabilized, mant-GDP, a nucleotide derivative, is reported to inhibit UCP1 (23). The remaining 10 compounds represent potential novel ligands of UCP1.

**Figure 3.**
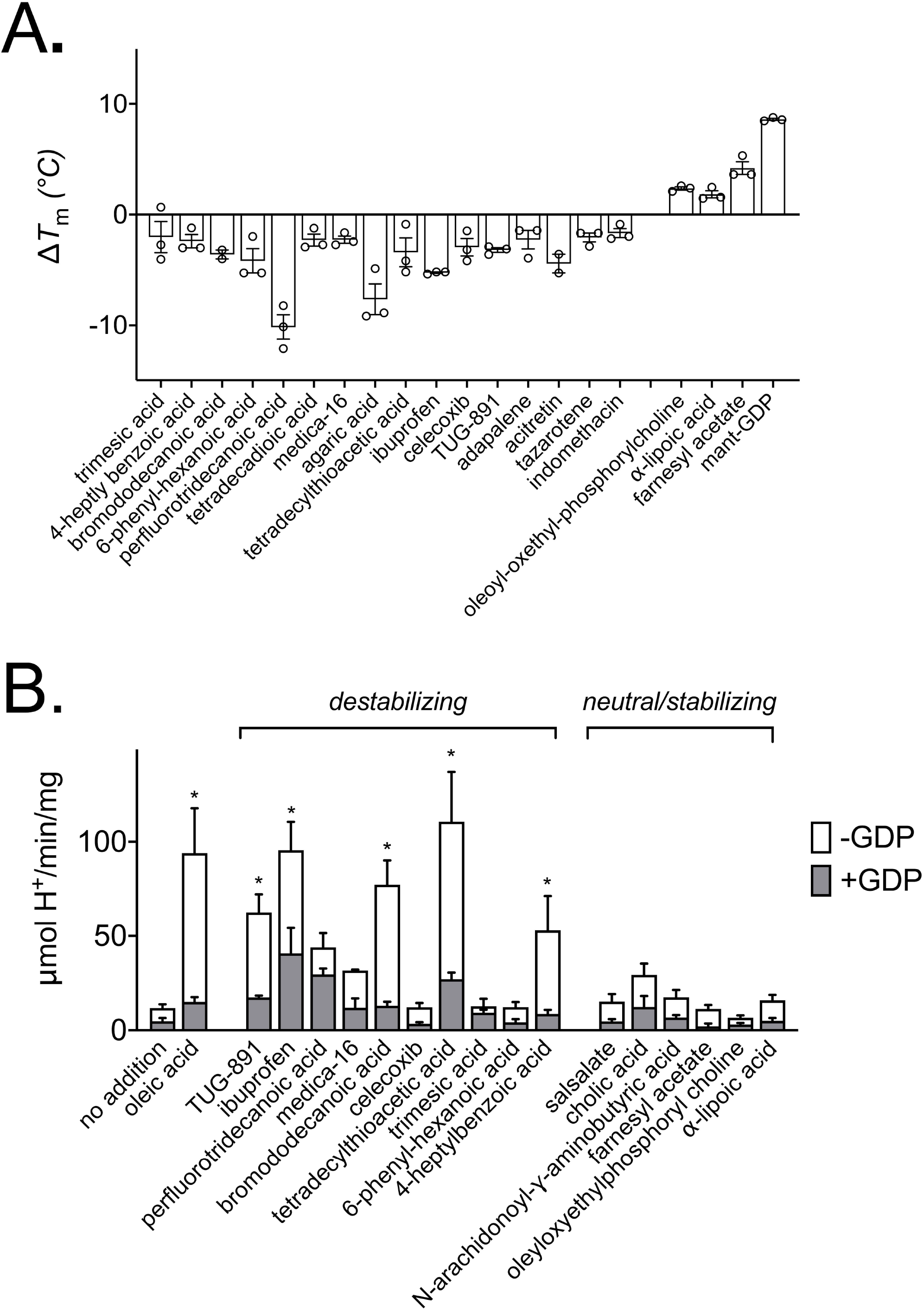
Novel activating ligands are found among compounds that destabilize UCP1 in stability screens. The relative thermal stability of purified UCP1 was determined as described for Fig. 1 (see Methods). A) Interacting ligands that induced a significant shift (Δ*T*_m_) in thermal stability of UCP1 following the screening of 72 notable compounds at 100 μM (see Fig EV2). Values are averages (±SD) of 3 independent experiments. (B) The proton leak activity (μmol H^+^/min/mg protein) by UCP1 in liposomes (± 1 mM GDP) induced by selected compounds following thermostability shift analysis. Activators of proton leak are found specifically among compounds that were identified through a destabilizing interaction with UCP1. Values are averages (±SEM) of 3-9 independent experiments. Statistical significance was determined by one-way ANOVA (\**p* < 0.05).

Compounds of interest were tested in UCP1 proteoliposome assays to determine their impact on proton leak activity. Strikingly, five of the destabilizing species tested proved to be activators of UCP1-mediated proton leak, as indicated by significantly increased proton conductance rates relative to controls, which were sensitive to GDP (Fig. 3B). In contrast, none of the compounds tested that either increased or had no effect on protein stability activated proton leak activity by UCP1. Nor did any of the non-nucleotide stabilizing ligands show significant inhibition of oleic acid-induced proton leak by UCP1 in follow up tests (Fig. EV3). These trends corroborate that the changes in UCP1 responsible for destabilization are likely an essential feature of activation. Importantly, the process identified three new compounds as direct activating ligands of the UCP1 protein: tetradecylthioacetic acid, a β-oxidation-resistant synthetic fatty acid that activates peroxisome proliferator-activated receptors (PPAR) and promotes mitochondrial fatty acid oxidation (34); TUG-891, a G protein coupled receptor 120 (GPR120) agonist reported to stimulate UCP1-dependent mitochondrial uncoupling, fat oxidation and weight loss in mice (35); and ibuprofen, a widely used non-steroidal anti-inflammatory drug and painkiller (36).

All of the UCP1-activating species observed were hydrophobic (logP >3.5) but amphipathic in nature, with a carboxylate head group that could be protonated at physiological pH to facilitate proton transport (p*K*_a_ >4), consistent with past observations (15, 22). However, the bulk hydrophobic region varied considerably from long chain fatty acids across activators (e.g. TUG-891), suggesting a lack of specific structural constraints. Computational alignment and analysis of verified activators revealed common matching features associated with the carboxylate headgroup for potential polar interactions but no specific features across the bulk region other than general hydrophobicity (see Fig. EV4A). The wider array of UCP1 ligands identified by thermostability analysis alone exhibited similar characteristics but with added diversity in the polar region and no strict requirement for a high-p*K*_a_ carboxylate group (see Fig. EV4B). These findings indicate that UCP1 has a particularly broad ligand specificity, which widens the scope of molecules with potential to activate the protein.

### Ibuprofen activates UCP1 activity in isolated brown adipose tissue mitochondria and HEK293 cells but not in immortalized brown adipocytes

Our screening process revealed that the licenced analgesic ibuprofen can activate isolated UCP1. Further investigations with the more soluble metabolic derivates indicated that only the parent molecule activated the protein in liposomes (Fig. EV5A), in line with the wider observed physiochemical trends. Tests with isolated mitochondria from mouse brown adipose tissue confirmed that ibuprofen could activate UCP1 in the natural membrane environment. Similar to palmitate, the drug decreased the membrane potential of succinate-energized mitochondria from WT mice in a GDP-sensitive manner, but not in mitochondria from UCP1 KO mice (see tests of safranin fluorescence quenching by mitochondria, Fig. 4A to C: with no activator, palmitate or ibuprofen addition, respectively). We tested the ability of ibuprofen to activate UCP1 in a cellular environment using both immortalized brown adipocytes from mice and a HEK293 cell UCP1-expression system (28). Whilst respiratory analysis did not indicate a stimulation of UCP1-dependent respiration in brown adipocytes (Fig. EV5 B and C), tests with HEK293 cells transfected with mouse UCP1 (MmUCP1) revealed a dose-dependent stimulation of non-phosphorylating respiration by ≥ 50 μM ibuprofen specific to UCP1-expressing cells (cf. oligomycin-insensitive oxygen consumption rates in UCP1-expressing vs empty vector control cells, Fig 4D-F). The drug was able to induce UCP1-dependent oxygen consumption similar to the TTNPB control, albeit requiring higher concentrations (~500 μM vs 15 μM TTNPB, cf. Fig. 4D-F). These findings provide a proof of principal for our process of identifying novel ligands that have the potential to activate UCP1-dependent energy expenditure in cells. The contrasting lack of response in cultured brown adipocytes suggests that other factors prevent the action of the drug on UCP1 in these cells, such as restricted uptake or a more rapid turnover to inactive variants. Circumventing these factors, e.g. via the development of suitable compound variants, therefore, is a promising new area for further research.

**Figure 4.**
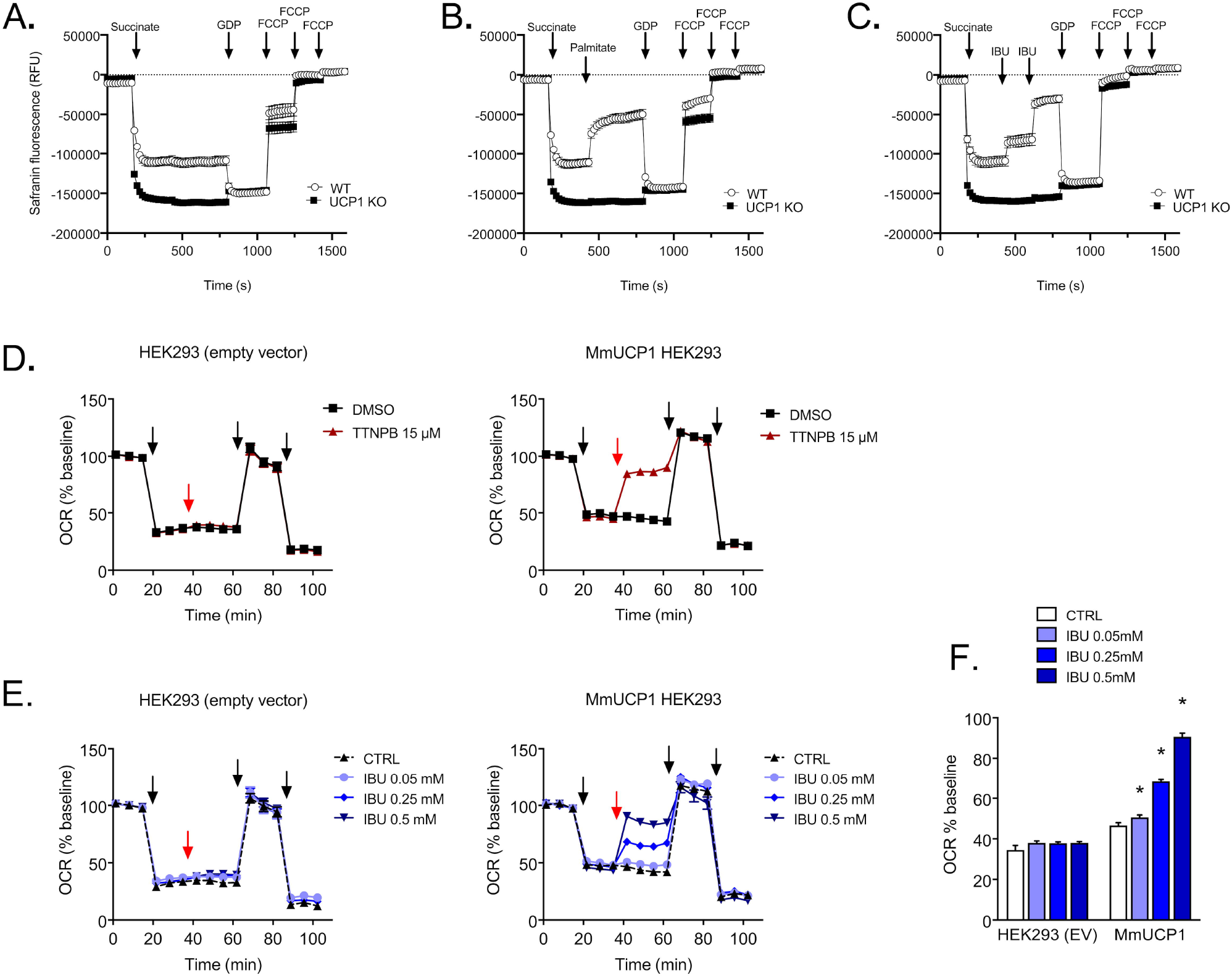
Ibuprofen stimulates UCP1 activity in mitochondria isolated from brown adipose tissue and in MmUCP1 (*Mus musculus*) transfected HEK293 cells. A-C: membrane potential measurements with safranin O for mitochondria isolated from wildtype (WT) or UCP1 KO mice, without (A) or with additions of 100 μM palmitate (B) or 2 × 250 μM ibuprofen (C) and other effectors, as indicated (see Materials and Methods). n=6 samples per group, measured on two independent days. Results are shown as mean ± SEM. (D and E) Oxygen consumption rate (OCR) of HEK293 cells transfected with an empty vector (EV) or mouse UCP1 (MmUCP1) upon treatment with TTNPB versus DMSO control (D) or different ibuprofen (IBU) concentrations versus buffer control (E). Arrows indicate injections, which are 1. Oligomycin, 2. TTNPB or ibuprofen (red arrow), 3. DNP and 4. rotenone + antimycin A (see Materials and Methods). BSA concentration of medium: 0.4%. (F) Summary of the OCR attributable to proton leak activity (oligomycin-insensitive respiration): treatment with ibuprofen stimulates UCP1 activity in a dose-dependent manner in MmUCP1 transfected HEK293 cells. Values (±SEM) are from 17 to 23 samples per group, measured on four independent days, with statistical significance determined by two-way ANOVA (\**p* < 0.05).

## Discussion

Integral membrane proteins like UCP1 are challenging to study. Hydrophobic ligands, such as fatty acids, can bind non-specifically and potentially act as harsh ionic detergents that denature proteins, hampering binding studies. Furthermore, UCP1 and other mitochondrial carriers are relatively unstable and can be extracted in detergent in an incorrectly folded form, leading to erroneous conclusions on function or biophysical properties (e.g. from common bacterial expression systems; see (24, 37–41) and references therein). Here, we have utilized a fluorescence-based protein thermostability assay tailored for membrane proteins to reveal details on the interaction of purified UCP1 with activators. The method reports on stability shifts related to the bonding changes of the protein itself, informing on interacting ligands and potential protein conformational shifts, avoiding issues of conventional binding studies. The methodology is emerging as a useful tool in resolving structural mechanisms and clarifying substrates in other carriers and transport proteins (20, 24, 27, 42, 43). Inherent to the technique is the monitoring of the folded state of the protein, which provides a robust indicator of protein sample integrity. Our experimental approach here not only reveals mechanistic information on how activators influence native UCP1, but also that the protein has a wide ligand specificity and may be directly targeted, independent of fatty acids, for increased energy expenditure.

Our thermostability measurements consistently demonstrated a destabilizing interaction of proton leak activators with purified UCP1, which matched the changes observed by equivalent alkyl sulfonate anions that do not facilitate proton leak but are transport substrates of the protein. The common destabilization profiles suggest the same bonding changes occur and that fatty acids and other activators interact as transport substrates, and that activator transport is fundamental in the UCP1 proton conductance mechanism. As such, our binding study observations support biochemical models of UCP1 proton leak where the protein specifically acts to transport the activating species (13, 15) over other models (see (9)). Accordingly, UCP1 exports bound fatty acid anions across the mitochondrial inner membrane, where they return to the matrix in a protonated form, either via UCP1 (*shuttling model* (15)) or by independently flipping directly across the membrane (*cycling model* (13)) to give a net proton conductance. In the former case, long chain fatty acid species are proposed to remain bound to UCP1 during transport in either direction, essentially chaperoning protons. Of the compounds that we observed to destabilize UCP1 that likely represent transport substrates, a subset proved to be activators. Notably, these all had p*K*_a_ values >4, consistent with the notion that it is the inherent ability of the molecule to be protonated in the relevant pH conditions that dictates whether or not a transported species facilitates proton leak, in line with previous evidence in support of these models (15, 22).

The distinct loss in stability from activator binding also provides clues to the structural process underlying UCP1 proton leak. The related mitochondrial ATP/ADP carrier exchanges nucleotides by a recently clarified structural mechanism, which is likely conserved across the mitochondrial carrier family of metabolite transporters (20). The protein cycles between a cytoplasmic (c-state) and matrix (m-state) conformation to alternate access of a substrate binding site within the central cavity of the protein to either side of the membrane for stepwise substrate transport. Substrate-binding induced state shifts occur through symmetrical movements in the three pseudo-symmetrically related domains of the carrier. Notably, deviations from symmetry occur in the central binding site to accommodate the non-symmetrical nature of the substrate. Similar to our observations here with UCP1, the purified ATP/ADP carrier was also *destabilized* by micromolar concentrations of ADP substrate, or by the m-state inhibitor bongkrekic acid, in similar conditions (24), consistent with ligand-induced transitioning to a less stable state of the transport cycle. The similar net bonding loss in UCP1 suggests that activating ‘substrates’ induce similar state shifts, and that activation uses the same core transport mechanism rather than a distinct asymmetrical process, as proposed (21, 44), where fatty acid anions translocate on the outside surface of the protein at the lipid-protein interface. UCP1 conserves all of the key structural elements of a conventional carrier mechanism, with no distinct asymmetrical features at the membrane-facing surface to facilitate specific interactions for ligand translocation (9), supporting this rationale. The region in UCP1 corresponding to the substrate binding site of common carriers (see (45)) has a triplet of arginine residues known to influence nucleotide binding (46, 47), which have potential to interact with the polar/charged group of activators as well, where hydrophobic residues in the central cavity could accommodate additional hydrophobic interactions. Notably, fatty acid activators and nucleotides have been reported to display competitive behaviour in regulating UCP1 (16).

Our screening approach successfully identified novel activating ligands among compounds that destabilized UCP1. The synthetic GPR120 agonist TUG-891 promotes brown adipose tissue thermogenic activity in mice through GPR120-signalling, but is also reported to induce UCP1-dependent oxygen consumption in isolated mitochondria, suggesting an additional mechanism (35). Our results corroborate these claims and demonstrate that TUG-891 is a direct activating ligand of the UCP1 protein, despite its structural deviation from conventional fatty acid activators. The positive health attributes observed in mice with this compound (increased fat oxidation, reduces body weight and fat mass) are at least in part likely due to the direct targeting of UCP1. The non-β-oxidizable fatty acid analogue, tetradecylthioacetic acid, which promotes mitochondrial fatty acid oxidation through PPAR activation (34), may also confer health benefits as a hypolipidemic agent via UCP1 targeting given our findings, though this notion requires verification. Surprisingly, we discovered that the cyclooxygenase inhibitor and common painkiller ibuprofen is an activating ligand of UCP1 as well, and could activate the protein in isolated brown fat mitochondria, HEK293 cells but not brown adipocytes. Ibuprofen may be sequestered and/or metabolized in brown adipocytes (e.g. by triglyceride incorporation, as can occur in white adipocytes (48)) that, if rapid enough, could prevent it from activating UCP1. Alternatively, the drug may not be taken up by these cells in contrast to kidney cell lines where uptake has been demonstrated to be mediated by an unknown transporter (49). Compound variants circumventing such barriers, therefore, are an attractive avenue for further investigation.

There is considerable research into methods to encourage brown fat proliferation and the browning of white adipose tissue, though in the absence of physiological stimuli, UCP1 must still be activated to benefit from the full glucose and triglyceride turnover capacity of the tissue (50). Pharmacological activation of upstream beta-adrenergic targets has been demonstrated in humans but lacks tissue specificity and is complicated by wider systemic effects (6, 51). As a specific, defining feature of brown fat, UCP1 is an attractive target for the therapeutic activation of thermogenic energy expenditure. Our assessment utilizing thermostability shift analysis provides an effective route to probe for UCP1-interacting molecules and reveals that the protein can be directly activated by conventional drug-like compounds with relatively wide structural specificity, which opens up the array of candidate molecules with potential to target it.

## Materials and Methods

### Materials

Dodecyl maltose neopentyl glycol (12MNG) detergent was obtained from Generon (Slough, UK). Lipids were obtained from Avanti Polar Lipids (AL, USA). Compounds were obtained from Cambridge Bioscience (Cambridge, UK) or Sigma-Aldrich (Merck Group). Unless otherwise stated, all other chemicals were obtained from Sigma-Aldrich (Merck Group) or Thermo Fisher Scientific.

### Tissue collection and isolation of mitochondria

Perirenal brown fat was collected from newborn lambs (10-40g per animal) that had died of natural causes at local Norfolk and Suffolk farms, and flash frozen in liquid nitrogen for storage until further use. Brown adipose tissue mitochondria were isolated following previously described methods (19), adapted from (52). Final mitochondrial samples (30-40 mg/mL protein) were flash frozen and stored in liquid nitrogen until further used.

Brown fat of WT and UCP1 KO mice ((53); age 8-10 weeks) was collected and put in ice cold buffer (250mM Sucrose, 10mM TES, 2mM EDTA and 1% defatted BSA (w/v), pH 7.2). The tissue was minced in buffer, on ice, with fine scissors and homogenized using a glass-Teflon homogenizer. The homogenate was filtered through nylon mesh and centrifuged (10 min, 8740 *g*, 4°C). Supernatant was discarded and the lipid remaining on the inside walls of the tube was removed using tissue paper. The pellet was resuspended in buffer (250mM Sucrose, 10mM TES, 1mM EGTA, 0.8% defatted BSA (w/v), pH 7.2) and centrifuged (10 min, 950 *g*, 4°C). The resulting supernatant was collected in a new tube and centrifuged (10 min, 8740 *g*, 4°C). The supernatant was discarded, and the pellet was resuspended in isolation buffer (100 mM KCl, 20 mM TES, 1 mM EGTA, pH 7.2). The solution was centrifuged (10 min, 8740 *g*, 4°C), and the final mitochondrial pellet was resuspended in a minimum volume of buffer (100mM KCl, 20mM TES, 1mM EGTA, pH 7.2). Mitochondrial protein content was determined using the Biuret method.

### Gene sequencing

The coding regions of the ovine UCP1 gene were verified by amplification of UCP1 exons from genomic DNA by polymerase chain reaction (PCR) and DNA sequencing.

Genomic DNA was isolated from ovine brown adipose tissue by phenol-chloroform extraction. ~300 mg of frozen tissue was ground, thawed and homogenised in 1.0 mL buffer (50 mM Tris-HCl pH 8, 25 mM EDTA, 400 mM NaCl, 1% SDS, 0.2 μg/μL proteinase K) and incubated at 65 **°**C for 2 hours. The sample was centrifuged (5 mins, ~16 000 g) and the supernatant recovered avoiding the pellet and top fat layer. 1/10 volumes of 5 M K-acetate (pH 7.5) was added and the sample incubated on ice for 30 mins, followed by centrifugation (10 mins, ~16 000 g). The supernatant was recovered in a fresh tube and vigorously mixed with an equal volume of phenol: chloroform: isoamyl alcohol (25:24:1). Following centrifugation (5 mins, ~16 000 g), the top layer was recovered (using a cut-off pipette tip) and a volume of chloroform added and vigorously mixed to the remaining bottom layer. Following re-centrifugation, the resulting top layer was recovered and pooled with the first top layer taken. Two volumes of ice-cold ethanol were mixed in and the sample incubated at −20 °C for 30 mins to precipitate the DNA. Following centrifugation (10 mins, 16 000 g), a DNA pellet was recovered and washed in 75% ethanol, re-centrifuged followed by a repeated wash. The final pellet was dried under a stream of nitrogen gas to remove excess water/ethanol and resuspended in nuclease-free water before quantification and further use.

Ovine genome sequence information ((54), gi957807226: 16832912-16839562, via NCBI: https://www.ncbi.nlm.nih.gov/gene/) was used to generate the primer pairs (below) to amplify the 6 exons of the ovine UCP1 gene by PCR. PCR products were verified and purified by standard agarose gel electrophoresis methods utilizing DNA sample clean up kits (Qiagen), and were sequenced using the same primers through commercial services (Source Bioscience, UK). The resulting data allowed the coding sequence for lamb UCP1 to be compiled based on established splicing (see Figure EV1), which highlighted a TAT codon (rather than TGT) encoding tyrosine at position 55 in the protein.

> <u>Exon 1</u> F1 AAGTCGGAGAGGACGGGTCT R1 TGGTAAAACCGAAGCCAGAGT <u>Exon 2</u> F2 AGATGTTGCTGAGAAGGCGAA R2 CCTGCAAGCTCTCCTGTGAT <u>Exon 3 and 4</u> F3 TGCACCAGCATTAGGGAAGT R3 ATACAGCTGCATCTTAGTTTTGTT <u>Exon 5</u> F4 GTCATGCCCTAAGGACAGAGG R4 TGCACATTCTAAGGCTGCTCA <u>Exon 6</u> F5 TCCTATACTCCATTCAGAAAGCAA R5 TGAATGTTTTGCTTTCCCTTCCT

#### Purification of native UCP1

UCP1 was purified using hydroxyapatite and thiopropyl sepharose chromatography media based on methods described in (19) and (55). 100 mg of thawed mitochondrial membranes were centrifuged (110,000 *g* for 20 min, °C) and the pellet resuspended and solubilized in 3% (v/v) Triton X-100 in purification medium (20 mM MOPS-NaOH pH 6.7, 20 mM Na_2_SO_4_, 0.16 mM EDTA) at 4°C for 20 min with gentle mixing. Insoluble material was pelleted by repeated ultracentrifugation and the supernatant recovered and loaded onto an hydroxyapatite column (7 g dry-weight resin, Biorad 130-0420, hydrated and packed in a disposable chromatography column, Biorad 732-1010, and pre-equilibrated in purification medium). Following column loading (0.7 mL/min), the flow was paused and the column incubated for 15 min at room temperature. Following addition of purification medium and re-starting the flow, ~11 mL of sample containing UCP1 was collected after the first 10 mL of eluate. The protein was immobilized and purified further by covalent chromatography. The sample was supplemented with 50 mM Tris pH 8.0 and 1 mM EDTA and applied to ~300 mg of thiopropyl Sepharose 6B (Sigma T8387; pre-hydrated in degassed water) in an empty PD-10 column, and rotated for 1 hour at 4 °C to allow UCP1 to bind to the resin. After settling, the column containing the sample was attached to a peristaltic pump and the resin carefully washed (~2.5 mL/min flow) with 30 mL wash buffer 1 (20 mM Tris pH 8, 0.5% 12MNG, ±0.5 mg/mL tetraoleoyl cardiolipin, 50 mM NaCl, 1 mM EDTA, followed by 30 mL wash buffer 2 (20 mM Tris pH 8, 0.005% 12MNG, ±0.005 mg/mL tetraoleoyl cardiolipin, 50 mM NaCl, 1 mM EDTA) to facilitate contaminant removal and detergent exchange. The column was detached from the pump, briefly centrifuged (500 g, 1 min) to remove excess liquid, and 1 mL of elution buffer (wash buffer 2 supplemented with 150 mM dithiothreitol) added to the resin before agitation in a cold room for 15 min. UCP1 was eluted by centrifugation (500 g, 1 min) and a further 0.8 mL of elution buffer added to the resin followed by repeated incubation. Following elution by centrifugation (2,000 g, 2 min), samples were pooled and the protein exchanged into sample buffer (20 mM Tris pH 8, 0.005% 12MNG, ±0.005 mg/mL tetraoleoyl cardiolipin) using a PD-10 column (17-0851-01; GE Healthcare) and protein quantified (BCA assay, Thermo Scientific) before flash freezing and storage in liquid nitrogen. Note, the supplementation of buffers with cardiolipin lipid, where indicated, was only carried out for UCP1 preparations used for liposome reconstitution experiments.

#### Protein thermostability shift measurements

Protein thermostability measurements were made using a fluorescence based assay suitable for membrane proteins (25) developed for enhanced and rapid use on a rotary qPCR machine with ‘melt’ analysis software (24). In the assay, CPM dye (7-Diethylamino-3-(4’-Maleimidylphenyl)-4-Methylcoumarin; Invitrogen D346) reacts with protein thiols as they become solvent exposed due to thermal denaturation to give a fluorescent adduct, which is used to monitor protein unfolding during an applied temperature ramp. In preparation for use, CPM dye stock (5 mg/mL; stored in DMSO at −80°C) was thawed and diluted 50-fold into assay buffer (20 mM Hepes pH 7.5, 0.1% 12MNG) and incubated in the dark for 10 to 15 min at room temperature. Per test, 2 μg of UCP1 protein was diluted into assay buffer, with test compounds added where required, to 45 μL in 200 μL thin-walled PCR tubes. Unless stated otherwise, 1 μL of test compound was added from a 50x stock or the appropriate solvent control (DMSO, ethanol or water). 5 μL of the diluted CPM solution was mixed into each test sample, which was subsequently incubated for 10-15 mins on ice. Samples were placed in a Rotor-Gene Q HRM 2-plex PCR cycler (36 position rotor) and subjected to a high resolution melt procedure (‘HRM’ channel: λ_ex_ 440-480 nm, λ_em_ 505-515 nm), with the instrument software set to give a required pre-measurement holding step (90 sec, beginning at 18 °C) followed by a temperature increase from 25 to 90 °C in 1 °C increments (‘wait between reading’ set to 4 sec), corresponding to a temperature ramp of 5.6 °C/min. Unless stated otherwise, protein unfolding profiles were analyzed using the instrument software, where the peak in the derivative provided a ‘melt temperature’ (*T*_m_) as a measure of relative protein stability. Typically, up to 18 tests were carried out per run, where *T*_m_ values gained in the presence of a given test compound/condition were subtracted from the corresponding ‘no compound’ solvent control to provide thermostability shift values (Δ*T*_m_) associated with compound-protein interaction.

#### Liposome reconstitution and proton flux measurements

UCP1 was reconstituted into liposomes based on methods described previously (19, 56), with adaptions for the use of the probe SPQ (6-Methoxy-N-(3-Sulfopropyl)Quinolinium, Invitrogen, M440) for proton transport measurements (13, 57). Ten milligrams of lipid (18:1 phosphatidylcholine supplemented with 5% 18:1 cardiolipin) was dried from storage in chloroform under a nitrogen stream, resolubilized in methanol, and re-dried to a smear in a 1.5-mL Eppendorf tube. The lipids were mixed with water and reagent stocks to form an emulsion and solubilized with 55 μL 25% (vol/vol) C_10_E_5_ detergent (Sigma 76436) on ice before addition of 20 μg of purified UCP1, to give a final ‘internal medium’ buffer composition of 100 mM K^+^ (phosphate salt, pH 7.5), 0.1 mM tetraethyl ammonium (TEA^+^; phosphate salt, pH 7.5), 30 mM TES (TEA^+^ salt, pH 7.5), 0.5 mM EDTA (TEA^+^ salt, pH 7.5) and 2 mM SPQ, with a final volume equivalent to 0.6 mL in the absence detergent. The detergent was removed through addition of 4 × 30 mg and a further 4 × 60 mg of adsorbent beads (Bio-Beads SM-2) in 20 min intervals with gentle mixing in a cold room. The resulting proteoliposomes were separated from the biobeads by using empty spin columns (Bio-Rad 732–6204) and treated with 40 mM methyl-β-cyclodextrin (Sigma C4555; 5-10 mins on ice) to sequester fatty acids, and exchanged into an external buffer [0.1 mM K^+^ (phosphate salt, pH 7.5), 100 mM TEA^+^ (phosphate salt, pH 7.5), 30 mM TES (TEA^+^ salt, pH 7.5), 0.5 mM EDTA (TEA^+^ salt, pH 7.5)] using a PD10 column. Following column loading (0.6 mL followed by 1.9 mL of buffer), the liposomes were collected in the first 1.2 mL of elution and stored on ice until further use.

Proton uptake into liposomes was tracked through changes in fluorescence of liposome-entrapped SPQ that is quenched specifically by the anionic component of the buffer (e.g. TES^-^), which changes in concentration in response to proton movement (see (57)). Measurements were made in a Cary Eclipse spectrofluorometer (λ_ex_ 334 nm, λ_em_ 443 nm) at 25 °C, where 75 μl of proteoliposomes were diluted to 500 μL in external buffer (see above) in a quartz cuvette, with 100 μM test compound, or 1 mM GDP, added from 100-fold or greater stocks solutions (i.e. in ≤ 5 μL volume), unless stated otherwise. Compound stocks were made up in water or ethanol (DMSO could not be used with the assay), where control traces included the equivalent volume of solvent alone. For each measurement, a signal was recorded for 40 seconds before a membrane potential was induced through the addition of the K^+^ ionophore valinomycin (2.5 μM) to drive proton uptake (added from a 250x stock in ethanol via a disposable inoculation loop). After a further 60 seconds, 1 μM CCCP was introduced in a similar manner to reveal the maximum proton uptake capacity of the system. Calibration of changes in signal to changes in internal proton concentration was achieved for each proteoliposome batch by diluting a 75 μL aliquot into internal buffer assay medium (rather than external buffer) and recording the signal changes associated with multiple 1 μL additions of 1 M H_2_SO_4_ in the presence 10 μM nigericin. The observed linear trends in Stern-Volmer plots (1/F vs δ[H^+^]) were modelled by linear regression to allow signal conversion in the corresponding progress curves as described in (57). The initial rate of change of proton concentration associated with UCP1 activity was estimated by fitting data 10 sec before to 59 sec after the addition of valinomycin to the exponential function ‘plateau and one phase association’ in GraphPad Prism software. The liposome internal volume was calculated by diluting a 75 μL aliquot into buffer with 0.2% Triton X-100 detergent to release entrapped SPQ, which was quantified by fluorescence using a standard curve after further SPQ additions of known concentration. Progress curves were corrected for internal volume to give the total number of protons moved per sample aliquot, per unit time for UCP1 rate estimations.

#### Ligand-based modelling perspective

Compounds were computationally aligned and analysed through ligand-based pharmacophore modelling software (Ligandscout). Once uploaded, the relevant compound structures were set to ionised acids and energy minimised. The set was clustered and conformers generated for each (‘maximum number of conformers’ set to 1000; ‘timeout’ to 1,200 sec, other settings to default). Pharmacophore models were subsequently generated, overnight where necessary, using the best fit and atom overlap with merged feature pharmacophore options to discern common features. For UCP1 proton leak activators, the following ten compounds were used: 4-heptylbenzoic acid; bromododecanoic acid; dodecanoic acid; ibuprofen; nonadecanoic acid; oleic acid; retinoic acid; tetradecylthioacetic acid; TTNPB; TUG-891. For the wider group of destabilizing ligands of UCP1, the following additional 11 compounds were also included: 1-octadecanesulfonic acid; 4-heptylbenzoic acid; 6-phenyl-hexanoic acid; acitretin; adapalene; agaric acid; celecoxib; medica-16; perfluorotridecanoic acid; tazarotene; tetradecadioic acid; undecane-1-sulfonate. In each case, the top outcome models were selected (scoring 0.81 and 0.77 for the activator and wider destabilizing ligand pharmacophore, respectively). Molecules were aligned by features; representative conformers are shown.

#### Cell culture methods and immortalization of mouse brown adipocytes

UCP1-expressing HEK293 cells were generated as described previously (58), and cultured in growth medium consisting of DMEM, high glucose (Gibco) with the addition of 10% Fetal Bovine Serum (Gibco), 1% Penicillin-Streptomycin solution (10,000 U/mL, Gibco) and 1% Geniticin selective Antibiotic (G418 Sulfate, 50 mg/mL, Gibco). Cells were trypsinized from culture dishes and seeded in appropriate amounts for experiments.

The isolation of the stromal vascular (SV) fraction from the interscapular brown adipose tissue (BAT) pad of WT and UCP1 KO mice (6-8 weeks old) was performed as follows. BAT pads were minced and then digested for 30 min at 37°C (DMEM/F12 plus glutamax, 0.15% (w/v) collagenase Type IV, 1% BSA). The homogenate was filtered through a 100 μm filter and rinsed with DMEM/ F12. Thereafter, the cells were centrifuged at 400 g for 10 min. The pellet was washed once more and centrifuged at 400 g for 10 min. The cells were re-suspended in growth media (DMEM/F12 plus glutamax, 10% FCS, 1% penicillin/streptomycin), plated and cultured. After 24 hrs, pre-adipocytes were immortalized using the SV40 large T-antigen. 48 hrs later, the cells were split, seeded into new flasks, allowed to grow to 60%–70% confluence and either further split or frozen into aliquots using freezing media (Growth media, 10% DMSO).

#### Extracellular flux measurements

HEK293 cells were seeded onto XF96 cell culture microplates (15k cells per well). The next day, the medium was changed to XF Assay medium (Agilent, 102353-100) with the addition of 10mM Glucose, 10mM Pyruvate and 0.4% defatted BSA (w/v) at pH 7.5 and incubated for an hour in an air incubator without CO_2_ at 37°C prior to the experiment. Three assay cycles (1 min mix, 2 min wait, 3 min measuring period) were used to determine basal respiration, followed by three assay cycles after oligomycin injection (4 μg/mL) to determine proton leak respiration. For the measurement of UCP1 activity, TTNPB (15 μM), ibuprofen (0.05, 0.25 and 0.5 mM) or a buffer control were used, followed by injection of chemical uncoupler (dinitrophenol, DNP at 100 μM) to determine maximal respiration and a final injection of rotenone (5 μM) and antimycin A (2 μM) to determine non-mitochondrial respiration. The measurements were performed in multiple wells on four independent experimental days (n= 17-23 wells per group).

For brown adipocytes, immortalized preadipocytes were plated onto XF96 cell culture micro plates (12K per well) and allowed to grow to 90%–100% confluence. At confluence, cell differentiation was started using differentiation cocktail tailored to brown adipocytes (Growth media, 5 μM dexamethasone, 0.5 mM IBMX, 1 μM rosiglitazone, 0.5 μg/ml insulin, 125 μM indomethacin, 1 nM T3) for 2 days, followed by a change to continuation medium (Growth media, 1 μM rosiglitazone, 0.5 μg/ml insulin, 1nM T3) for 2 days, and a change to differentiation medium (Growth media, 0.5 μg/ml Insulin, 1 nM T3) for 2 days. On day six of differentiation the cells were used for cellular respiration assays. Adipocytes were switched to XF Assay medium (Agilent, 102353-100) with the addition of 20 mM glucose, 2 mM glutamine and 0.4% defatted BSA (w/v) at pH 7.2. The cells were incubated for 10 min in an air incubator without CO_2_ at 37°C. Four assay cycles (2 min mix, 0 min waiting and 2 min measuring period) were used to determine basal respiration, followed by three assay cycles after oligomycin injection (5 μg/mL) to determine proton leak respiration. For the measurement of UCP1 activation, isoproterenol (0.5 μM), ibuprofen (0.05, 0.25 and 0.5 mM) or a buffer control were used, followed by dinitrophenol (DNP; 150 μM) injection to determine maximal respiration and a final injection of rotenone (5 μM) and antimycin A (2 μM) to determine non-mitochondrial respiration. These measurements were performed in multiple wells on three independent experimental days (n = 9 wells per group).

#### Mitochondrial membrane potential measurements

Measurements of Safranin O fluorescence were done in a 96-well plate with the Clariostar (λ_ex_ 533-15 nm, λ_em_ 576-20 nm, bottom optic, gain 1670, focal height of 5.4mm, cycle time of 18 s and orbital shaking (30 s at 300 rpm) before the first measurement). Per well, 40μg of mitochondria were used along with 200μL of pre-warmed KHE buffer (50mM KCl, 5mM TES, 2mM MgCl_2_ 6*H_2_O, 4mM KH_2_PO_4_, 1mM EGTA, 0,8% defatted BSA (w/v), pH 7.2) mixed with safranin O (10 μM), nigericin (100 nM), oligomycin (1 μg/mL) and rotenone (4μM). A baseline measurement was taken for 10 cycles. After, succinate (5 mM) was added to each well, and the measurement continued for 15 cycles. For the measurement of UCP1 activity, palmitate (100μM) and ibuprofen (250 μM) were used, followed by injection of GDP (1 mM). Lastly, FCCP was titrated in all wells until maximum fluorescence was reached (steps of 6 μM, 3 μM and 3 μM), with 10 measuring cycles after each addition. The measurements were performed in multiple wells on two independent experimental days (n= 6 wells per group).

#### Western blotting

During cultivation, WT and UCP1 KO immortalized brown adipocytes were collected for protein extraction using RIPA-based lysis buffer (150 mM NaCl, 1% IGEPAL CA-630, 0.5% sodium deoxycholate, 0.1% SDS, 50 mM Tris, pH 8.0) with a phosphatase/protease inhibitor. Differentiated adipocytes (day 6) were scraped off the 6-well plate and incubated on ice for 30°C. Thereafter, samples were centrifuged for 30 min at 18.000 *g* at 4°C, and the supernatant was collected. Protein concentration was quantified with a Bradford assay (Sigma). The following primary antibodies were used: UCP1 (1:2000, Abcam, ab23841) and α Tubulin Antibody (1:2000, Santa Cruz, sc-23948). The following Horseradish-peroxidase-conjugated secondary antibodies were used: Goat Anti-Rabbit IgG H&L (HRP) (1:10.000, Abcam, ab6721), Anti-β-Actin Antibody (1:2000, Santa Cruz, sc-47778) and Goat Anti-mouse IgG-HRP (1:10.000, Santa Cruz, sc-2005). The immunoblot was visualized with chemiluminescence (Clarity™ Western ECL Substrate, 1705060, BioRad).

#### Statistical tests

Statistical analyses were performed using GraphPad Prism software by one-way ANOVA with Dunnett’s post-hoc comparison test where *p* values < 0.05 were considered significant. For thermostability shift values (Δ*T*_m_), individual *T*_m_ control values were subtracted from *T*_m_ control averages to give the control Δ*T*_m_ zero value with standard deviation for comparison with test values. For HEK293 cell extracellular flux measurements, statistical significance was determined by two-way ANOVA and Holm-Sidak post-hoc analysis (\**p* < 0.05).

## Acknowledgements

We would like to thank local Norfolk and Suffolk farms for access to new-born lambs that died of natural causes during the lambing season. This work was supported a PhD studentship award (R.C.) from Norwich Medical School and the Faculty of Medicine and Health, University of East Anglia, UK, and by funding from the UK Biological and Biotechnological Sciences Research Council (BB/S00940X/1), Novo Nordisk Research Foundation (0059646 to M.J.) and Swedish Research Council (2018-02150 to S.K).

## Abbreviations

UCP1: Uncoupling protein 1
12MNG: dodecyl maltose neopentyl glycol (2,2-didecylpropane-1,3-bis-β-D-maltopyranoside)
OLPA: oleoyl-L-α-lysophosphatidic acid
TTNPB: 4-[(E)-2-(5,6,7,8-Tetrahydro-5,5,8,8-tetramethyl-2-naphthalenyl)-1-propenyl]benzoic acid
C7: heptanoic acid
C12: dodecanoic acid
C19: nonadecanoic acid
C6-S: hexane sulfonate
C11-S: undecane sulfonate
C18-S: octadecane sulfonate

## Expanded View Figures Legends

**Figure EV1.**
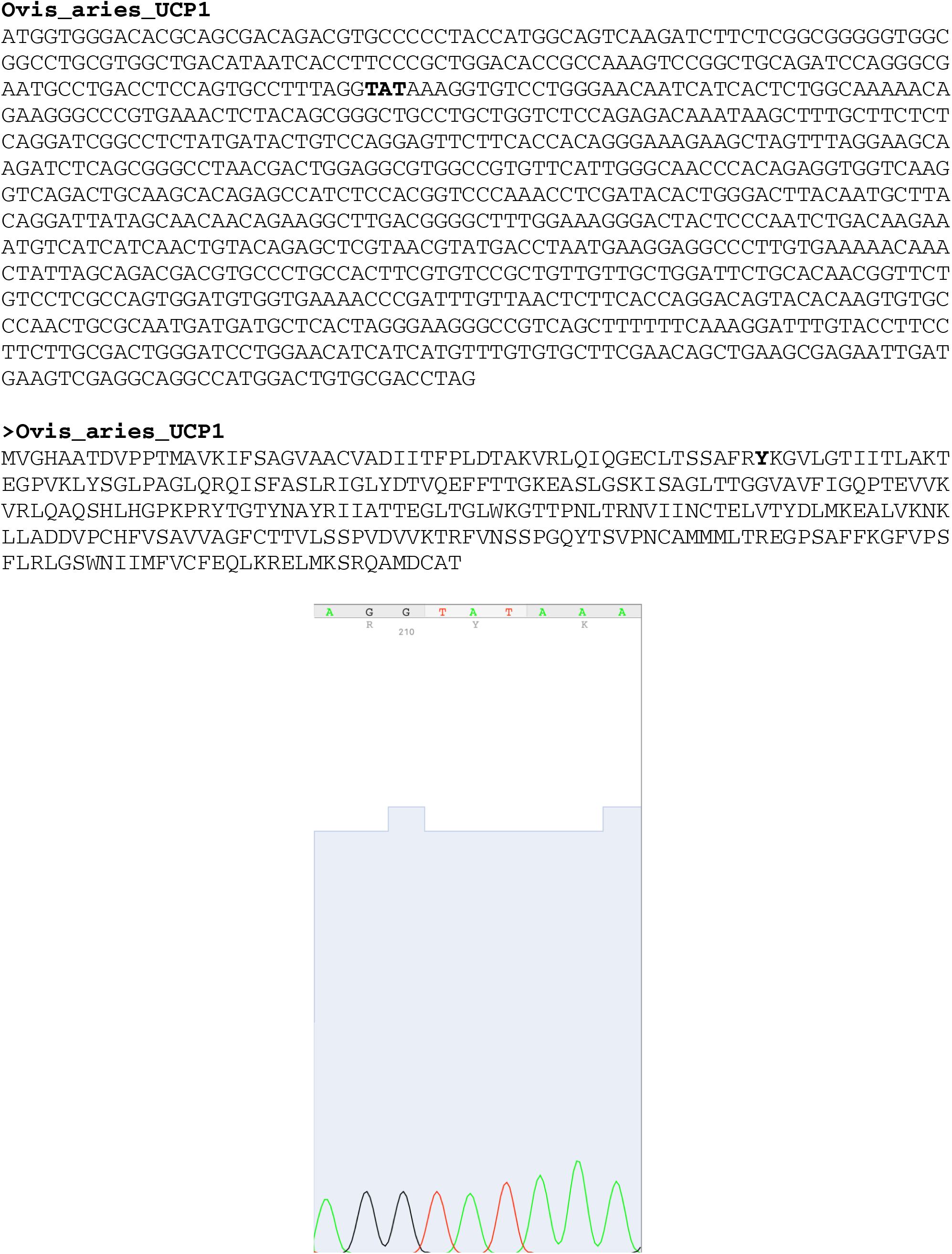
The gene coding sequence (top) and corresponding amino acid sequence (middle) of ovine UCP1 determined by exon sequencing as described in the methods section. Tyrosine 55 (corresponding to the TAT codon), in bold, matches the tyrosine conservation at this position in UCP1 in other species, but contrasts with a cysteine (TGT codon) at this position in other database submissions for the ovine protein (e.g. (26); see (19) for example amino acid sequence alignment). The raw sequencing data for the tyrosine 55 codon is also given (bottom)

**Figure EV2.**
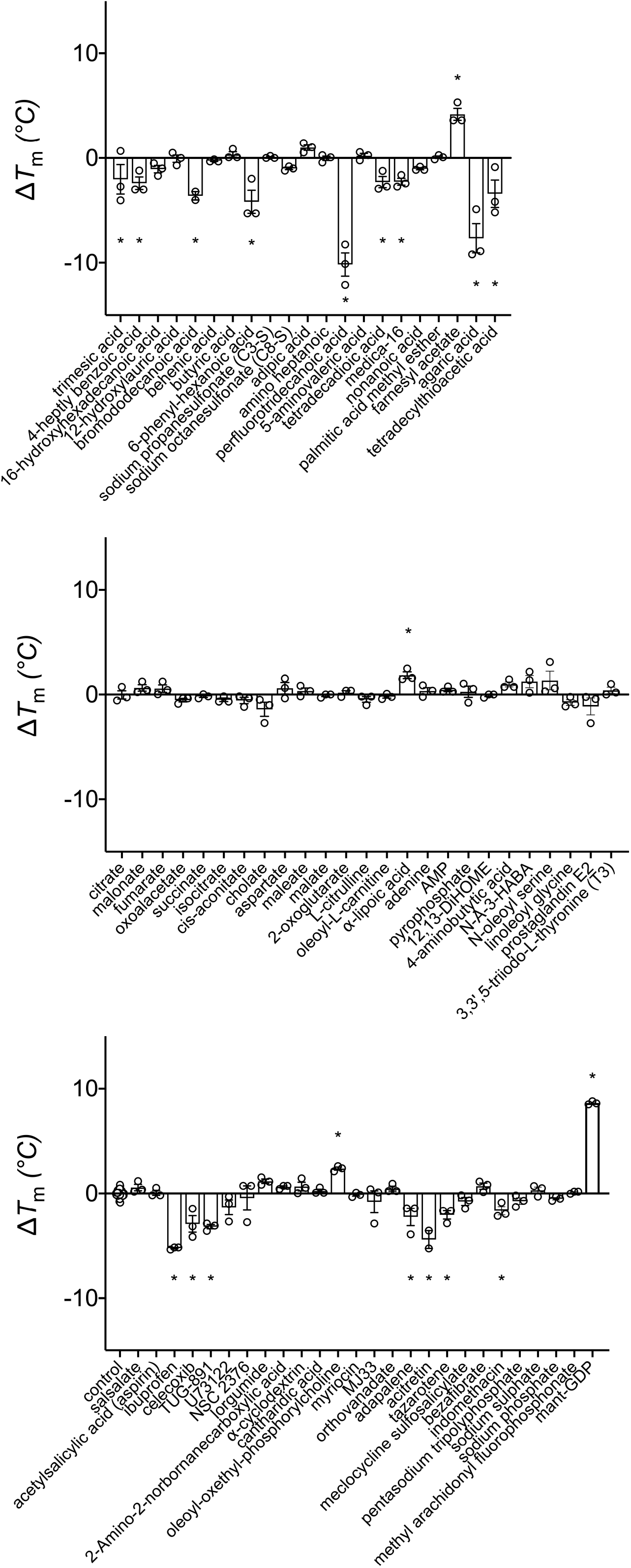
The identification of UCP1 ligands by screening compounds for shifts in protein thermostability. The relative thermal stability of purified UCP1 was determined as described in Methods. 72 compounds were tested at 100 μM to identify interacting ligands that induced a significant shift (Δ*T*_m_; condition minus control) in thermal stability of UCP1. Values shown are averages (±SEM) of 3 independent experiments. Statistical significance was determined by one-way ANOVA tests (\**p* < 0.05).

**Figure EV3.**
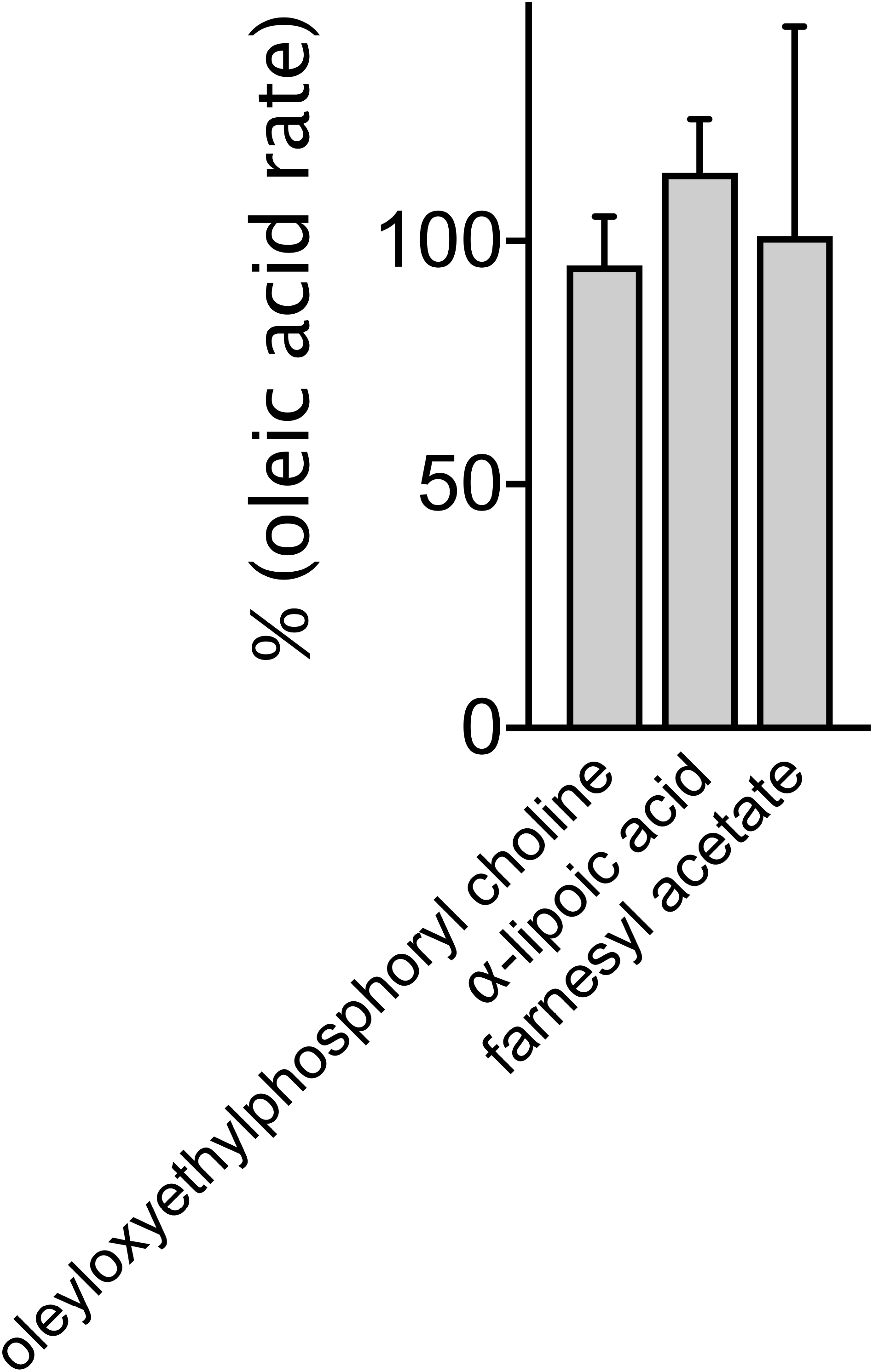
The effect of stabilizing compounds on oleic acid-activated rates of proton leak activity. Rates were measured in the presence of 100 μM oleic acid and 100 μM of the indicated compound. Values are expressed as a percentage of the rate obtained in the presence of oleic acid alone and are averages (±SD) of 3-4 independent experiments.

**Figure EV4.**
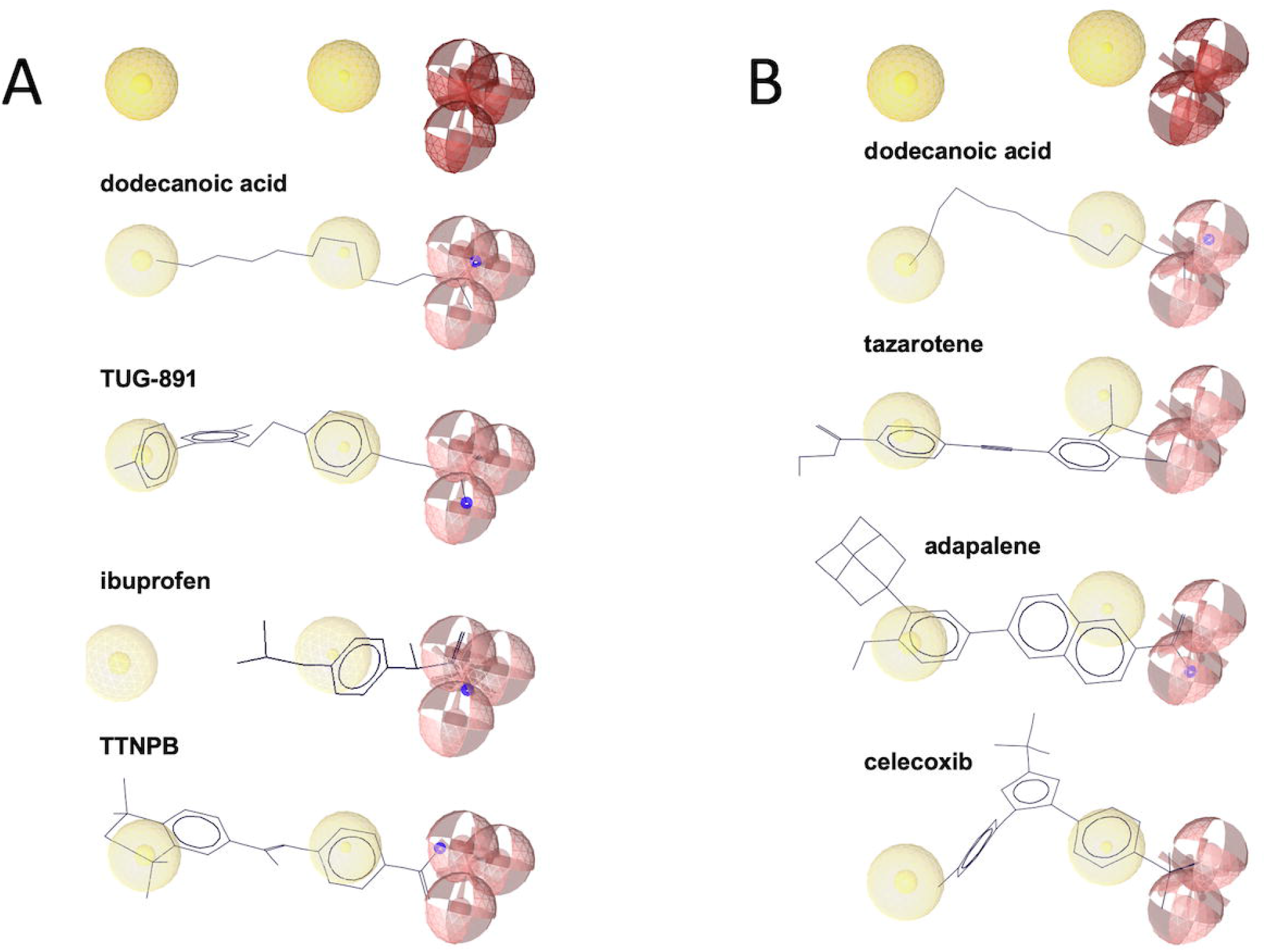
*In silico* pharmacophore modelling for common features in proton leak activators alone (A) or with the inclusion of wider destabilizing ligands (B) of UCP1. Models were generated as described in the Materials and Methods. In each model (top), the following common matching features are shown: hydrogen bond acceptor (red spheres), negative ionizable area (red star) and hydrophobic interactions (yellow spheres). Below each model alignments of example molecules from each group are given (negative charges are shown in blue). Ten activators were used alone or as part of a wider group of 21 destabilizing ligands in each case.

**Figure EV5.**
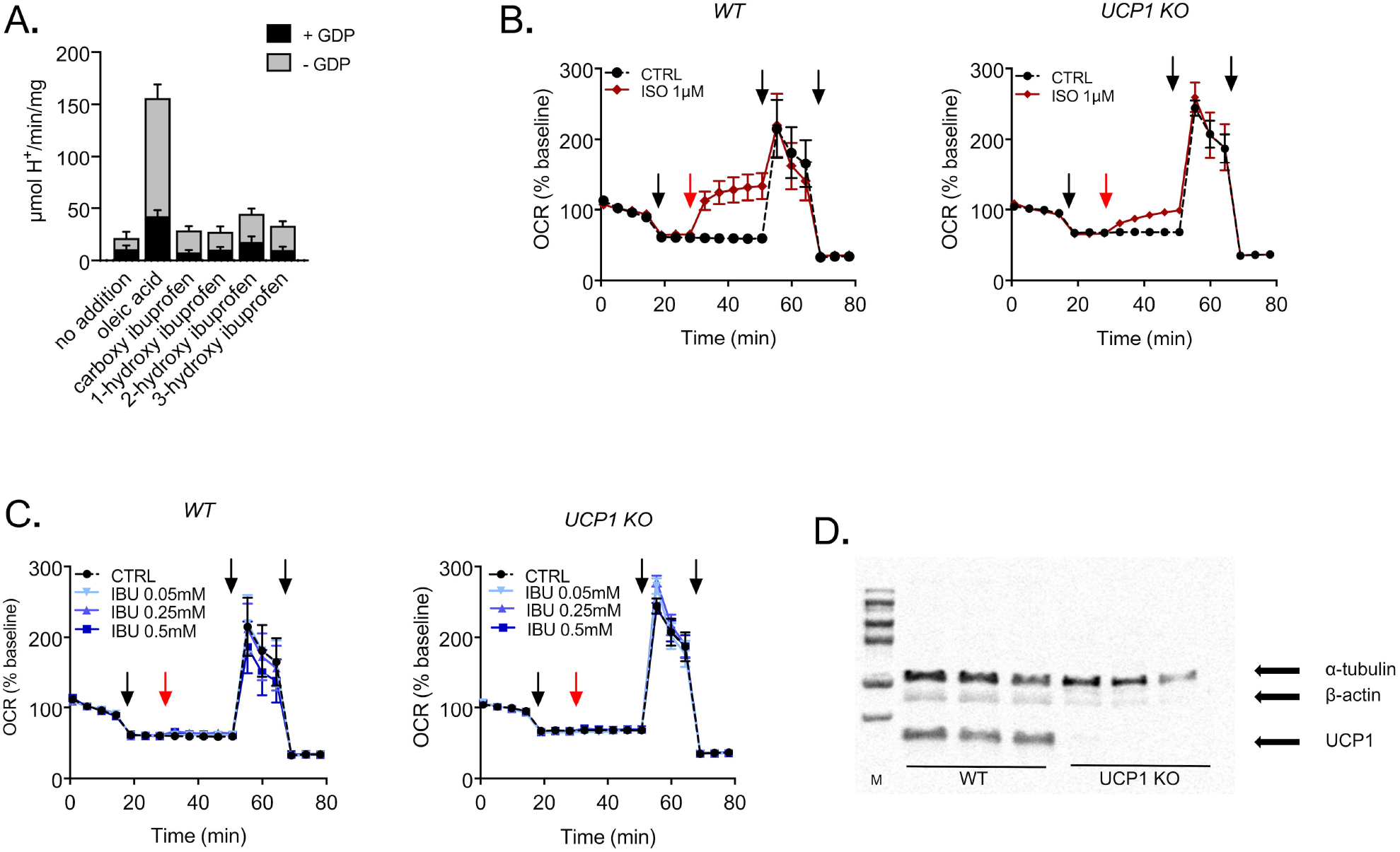
The effect of ibuprofen derivates on UCP1 activity in liposomes, and ibuprofen on UCP1 activity in immortalized brown adipocyte (iBAT) cell cultures. (A) The metabolic derivates of ibuprofen do not stimulate proton leak activity by ovine UCP1 in liposomes (± 1mM GDP). Values are averages (±SEM) of three to six independent experiments. B and C: ibuprofen does not stimulate UCP1-dependent respiration in iBAT cells. Oxygen consumption rate (OCR) of WT and UCP1 KO iBAT cells upon treatment with isoproterenol versus buffer medium CTRL (B), or ibuprofen versus buffer medium CTRL (C), at the concentrations indicated. n=9 samples per group, measured on three independent days. Arrows indicate injections, which are 1. Oligomycin, 2. Isoproterenol or ibuprofen (red arrow), 3. DNP and 4. Rotenone + Antimycin A (see Materials and Methods). BSA concentration of medium: 0.4%. Values (±SEM) are from 9 samples per group, measured on 3 independent days. D: UCP1 protein expression in WT iBAT cells versus no UCP1 protein expression in UCP1 KO iBAT cells.

## Notes

The authors declare no conflict of interest

### Competing Interest Statement

The authors have declared no competing interest.

## Bibliography

1. B. Cannon, J. Nedergaard, Brown adipose tissue: function and physiological significance. Physiol Rev 84, 277–359 (2004).

2. T. Yoneshiro et al., Age-related decrease in cold-activated brown adipose tissue and accumulation of body fat in healthy humans. Obesity (Silver Spring) 19, 1755–1760 (2011).

3. A. M. Cypess et al., Identification and importance of brown adipose tissue in adult humans. N Engl J Med 360, 1509–1517 (2009).

4. J. Nedergaard, T. Bengtsson, B. Cannon, Unexpected evidence for active brown adipose tissue in adult humans. Am J Physiol Endocrinol Metab 293, E444–452 (2007).

5. M. Chondronikola et al., Brown adipose tissue improves whole-body glucose homeostasis and insulin sensitivity in humans. Diabetes 63, 4089–4099 (2014).

6. A. M. Cypess et al., Activation of human brown adipose tissue by a beta3-adrenergic receptor agonist. Cell Metab 21, 33–38 (2015).

7. M. Saito et al., High incidence of metabolically active brown adipose tissue in healthy adult humans: effects of cold exposure and adiposity. Diabetes 58, 1526–1531 (2009).

8. M. Chondronikola et al., Brown Adipose Tissue Activation Is Linked to Distinct Systemic Effects on Lipid Metabolism in Humans. Cell Metab 23, 1200–1206 (2016).

9. P. G. Crichton, Y. Lee, E. R. Kunji, The molecular features of uncoupling protein 1 support a conventional mitochondrial carrier-like mechanism. Biochimie 134, 35–50 (2017).

10. R. M. Locke, E. Rial, I. D. Scott, D. G. Nicholls, Fatty acids as acute regulators of the proton conductance of hamster brown-fat mitochondria. Eur J Biochem 129, 373–380 (1982).

11. A. M. Singh et al., Human beige adipocytes for drug discovery and cell therapy in metabolic diseases. Nat Commun 11, 2758 (2020).

12. M. Klingenberg, S. G. Huang, Structure and function of the uncoupling protein from brown adipose tissue. Biochim Biophys Acta 1415, 271–296 (1999).

13. K. D. Garlid, D. E. Orosz, M. Modriansky, S. Vassanelli, P. Jezek, On the mechanism of fatty acid-induced proton transport by mitochondrial uncoupling protein. J Biol Chem 271, 2615–2620 (1996).

14. K. D. Garlid, M. Jaburek, P. Jezek, The mechanism of proton transport mediated by mitochondrial uncoupling proteins. FEBS Lett 438, 10–14 (1998).

15. A. Fedorenko, P. V. Lishko, Y. Kirichok, Mechanism of fatty-acid-dependent UCP1 uncoupling in brown fat mitochondria. Cell 151, 400–413 (2012).

16. I. G. Shabalina, A. Jacobsson, B. Cannon, J. Nedergaard, Native UCP1 displays simple competitive kinetics between the regulators purine nucleotides and fatty acids. J Biol Chem 279, 38236–38248 (2004).

17. E. Pebay-Peyroula et al., Structure of mitochondrial ADP/ATP carrier in complex with carboxyatractyloside. Nature 426, 39–44 (2003).

18. H. Aquila, T. A. Link, M. Klingenberg, Solute carriers involved in energy transfer of mitochondria form a homologous protein family. FEBS Lett 212, 1–9 (1987).

19. Y. Lee, C. Willers, E. R. Kunji, P. G. Crichton, Uncoupling protein 1 binds one nucleotide per monomer and is stabilized by tightly bound cardiolipin. Proc Natl Acad Sci U S A 112, 6973–6978 (2015).

20. J. J. Ruprecht et al., The Molecular Mechanism of Transport by the Mitochondrial ADP/ATP Carrier. Cell 176, 435–447.e415 (2019).

21. P. Jezek, M. Jaburek, K. D. Garlid, Channel character of uncoupling protein-mediated transport. FEBS Lett 584, 2135–2141 (2010).

22. P. Jezek, M. Modriansky, K. D. Garlid, A structure-activity study of fatty acid interaction with mitochondrial uncoupling protein. FEBS Lett 408, 166–170 (1997).

23. A. S. Divakaruni, D. M. Humphrey, M. D. Brand, Fatty acids change the conformation of uncoupling protein 1 (UCP1). J Biol Chem 287, 36845–36853 (2012).

24. P. G. Crichton et al., Trends in thermostability provide information on the nature of substrate, inhibitor, and lipid interactions with mitochondrial carriers. J Biol Chem 290, 8206–8217 (2015).

25. A. I. Alexandrov, M. Mileni, E. Y. Chien, M. A. Hanson, R. C. Stevens, Microscale fluorescent thermal stability assay for membrane proteins. Structure 16, 351–359 (2008).

26. Y. N. Yuan, W. Z. Liu, J. H. Liu, L. Y. Qiao, J. L. Wu, Cloning and ontogenetic expression of the uncoupling protein 1 gene UCP1 in sheep. J Appl Genet 53, 203–212 (2012).

27. H. Majd et al., Screening of candidate substrates and coupling ions of transporters by thermostability shift assays. Elife 7 (2018).

28. S. Keipert, M. Jastroch, Brite/beige fat and UCP1 - is it thermogenesis? Biochim Biophys Acta 1837, 1075–1082 (2014).

29. E. Rial et al., Retinoids activate proton transport by the uncoupling proteins UCP1 and UCP2. Embo j 18, 5827–5833 (1999).

30. M. Klingenberg, Ligand-protein interaction in biomembrane carriers. The induced transition fit of transport catalysis. Biochemistry 44, 8563–8570 (2005).

31. P. Jezek, K. D. Garlid, New substrates and competitive inhibitors of the Cltranslocating pathway of the uncoupling protein of brown adipose tissue mitochondria. J Biol Chem 265, 19303–19311 (1990).

32. E. Winkler, M. Klingenberg, Effect of fatty acids on H+ transport activity of the reconstituted uncoupling protein. J Biol Chem 269, 2508–2515 (1994).

33. I. G. Shabalina, E. C. Backlund, J. Bar-Tana, B. Cannon, J. Nedergaard, Within brown-fat cells, UCP1-mediated fatty acid-induced uncoupling is independent of fatty acid metabolism. Biochim Biophys Acta 1777, 642–650 (2008).

34. B. Bjørndal, L. Burri, V. Staalesen, J. Skorve, R. K. Berge, Different adipose depots: their role in the development of metabolic syndrome and mitochondrial response to hypolipidemic agents. J Obes 2011, 490650 (2011).

35. M. Schilperoort et al., The GPR120 agonist TUG-891 promotes metabolic health by stimulating mitochondrial respiration in brown fat. EMBO Mol Med 10 (2018).

36. K. D. Rainsford, Ibuprofen: pharmacology, efficacy and safety. Inflammopharmacology 17, 275–342 (2009).

37. M. S. King, P. G. Crichton, J. J. Ruprecht, E. R. S. Kunji, Concerns with yeast mitochondrial ADP/ATP carrier’s integrity in DPC. Nat Struct Mol Biol 25, 747–749 (2018).

38. M. S. Piel, S. Masscheleyn, F. Bouillaud, K. Moncoq, B. Miroux, Structural models of mitochondrial uncoupling proteins obtained in DPC micelles are not functionally relevant. Febs j 10.1111/febs.15629 (2020).

39. V. Kurauskas et al., How Detergent Impacts Membrane Proteins: Atomic-Level Views of Mitochondrial Carriers in Dodecylphosphocholine. J Phys Chem Lett 9, 933–938 (2018).

40. F. Dehez, P. Schanda, M. S. King, E. R. S. Kunji, C. Chipot, Mitochondrial ADP/ATP Carrier in Dodecylphosphocholine Binds Cardiolipins with Non-native Affinity. Biophys J 113, 2311–2315 (2017).

41. M. Zoonens et al., Dangerous liaisons between detergents and membrane proteins. The case of mitochondrial uncoupling protein 2. J Am Chem Soc 135, 15174–15182 (2013).

42. M. S. King, M. Kerr, P. G. Crichton, R. Springett, E. R. Kunji, Formation of a cytoplasmic salt bridge network in the matrix state is a fundamental step in the transport mechanism of the mitochondrial ADP/ATP carrier. Biochim Biophys Acta 1857, 14–22 (2016).

43. S. P. Harborne, M. S. King, P. G. Crichton, E. R. Kunji, Calcium regulation of the human mitochondrial ATP-Mg/Pi carrier SLC25A24 uses a locking pin mechanism. Sci Rep 7, 45383 (2017).

44. P. Ježek, M. Jabůrek, R. K. Porter, Uncoupling mechanism and redox regulation of mitochondrial uncoupling protein 1 (UCP1). Biochim Biophys Acta Bioenerg 1860, 259–269 (2019).

45. A. J. Robinson, E. R. Kunji, Mitochondrial carriers in the cytoplasmic state have a common substrate binding site. Proc Natl Acad Sci U S A 103, 2617–2622 (2006).

46. M. Modriansky, D. L. Murdza-Inglis, H. V. Patel, K. B. Freeman, K. D. Garlid, Identification by site-directed mutagenesis of three arginines in uncoupling protein that are essential for nucleotide binding and inhibition. J Biol Chem 272, 24759–24762 (1997).

47. M. Klingenberg, K. S. Echtay, Uncoupling proteins: the issues from a biochemist point of view. Biochim Biophys Acta 1504, 128–143 (2001).

48. S. Vickery, P. F. Dodds, Incorporation of xenobiotic carboxylic acids into lipids by cultured 3T3-L1 adipocytes. Xenobiotica 34, 1025–1042 (2004).

49. C. U. Nielsen et al., A Transporter of Ibuprofen is Upregulated in MDCK I Cells under Hyperosmotic Culture Conditions. Mol Pharm 13, 3119–3129 (2016).

50. J. Nedergaard, B. Cannon, The changed metabolic world with human brown adipose tissue: therapeutic visions. Cell Metab 11, 268–272 (2010).

51. A. S. Baskin et al., Regulation of Human Adipose Tissue Activation, Gallbladder Size, and Bile Acid Metabolism by a β3-Adrenergic Receptor Agonist. Diabetes 67, 2113–2125 (2018).

52. B. Cannon, O. Lindberg, Mitochondria from brown adipose tissue: isolation and properties. Methods Enzymol 55, 65–78 (1979).

53. S. Keipert et al., Long-Term Cold Adaptation Does Not Require FGF21 or UCP1. Cell Metab 26, 437–446.e435 (2017).

54. A. L. Archibald et al., The sheep genome reference sequence: a work in progress. Anim Genet 41, 449–453 (2010).

55. C. S. Lin, M. Klingenberg, Isolation of the uncoupling protein from brown adipose tissue mitochondria. FEBS Lett 113, 299–303 (1980).

56. K. S. Echtay, M. Bienengraeber, M. Klingenberg, Mutagenesis of the uncoupling protein of brown adipose tissue. Neutralization Of E190 largely abolishes pH control of nucleotide binding. Biochemistry 36, 8253–8260 (1997).

57. D. E. Orosz, K. D. Garlid, A sensitive new fluorescence assay for measuring proton transport across liposomal membranes. Anal Biochem 210, 7–15 (1993).

58. M. Jastroch, Expression of uncoupling proteins in a mammalian cell culture system (HEK293) and assessment of their protein function. Methods Mol Biol 810, 153–164 (2012).

